# Unfolded Protein Response signaling restricts PTI and limits salicylic acid-dependent cell death under ER stress

**DOI:** 10.64898/2026.10.02.755983

**Authors:** Veronika Cehulová, Anzhela Antonova, Hana Leontovyčová, Lenka Burketová, Amélie Ducloy, Marianne Azzopardi, Jean-Luc Cacas, Tetiana Kalachova

## Abstract

The Unfolded Protein Response (UPR) maintains endoplasmic reticulum (ER) homeostasis during proteotoxic stress while also contributing to plant immunity. However, the functional relationship between UPR signaling, pathogen-associated molecular pattern (PAMP)-triggered immunity (PTI) and salicylic acid (SA) remains poorly understood. Here, we investigated this interplay using Arabidopsis mutants impaired in the two major UPR branches. Mutants defective in the INOSITOL-REQUIRING ENZYME 1 (IRE1)-bZIP60 branch displayed enhanced resistance to *Pseudomonas syringae* pv. tomato *DC3000* and to the type III secretion-deficient strain hrcC^−^, indicating potentiation of PTI. Consistently, the *ire1a ire1b* double mutant exhibited increased flg22-induced callose deposition, whereas reactive oxygen species production was unaffected. Despite enhanced basal resistance, all UPR mutants retained full responsiveness to benzothiadiazole, demonstrating that downstream SA signaling remains functional independently of UPR integrity. Conversely, acute ER stress induced by tunicamycin promoted SA accumulation and extensive leaf necrosis. Reverse genetics revealed that SA-deficient mutants were largely protected from ER stress-induced cell death, whereas mutants over-accumulating SA exhibited enhanced susceptibility, demonstrating that endogenous SA is required for the full execution of this response. Analysis of an independent transcriptomic dataset further showed that loss of *IRE1* de-represses genes involved in SA biosynthesis, PTI and defense under ER stress. Together, our results identify IRE1 as a key regulator coordinating ER homeostasis with immune signaling by limiting SA-dependent defense activation during ER stress, while simultaneously preventing excessive cell death.

**Highlights:**

- UPR deficiency primes PAMP-triggered immunity and increases resistance to bacterial pathogens.
- The UPR acts as a brake on PTI, inverting the view of the UPR as a simple positive regulator of immunity.
- Acute ER stress drives salicylic acid accumulation
- ER stress-induced cell death is dependent on SA
- IRE1, bZIP60 and bZIP28 are the principal UPR actors mitigating acute tunicamycin-induced ER stress.
- BTH-induced resistance is independent of the UPR, separating induced resistance from ER stress control.

## 1. Introduction

Plants are continuously exposed to fluctuating environmental conditions and must rapidly adapt to a wide range of biotic and abiotic stresses. To ensure survival under adverse conditions, plant cells have evolved sophisticated regulatory networks that coordinate stress perception, signal transduction, and metabolic reprogramming. Stress adaptation often requires the rapid synthesis of new proteins, which can substantially increase the load imposed on the endoplasmic reticulum (ER) as the major site of secreted protein folding and maturation in the endomembrane system. In addition, environmental stresses may directly perturb ER homeostasis, affecting protein folding efficiency and quality control mechanisms. When the protein folding and degradation capacity of the ER becomes insufficient, misfolded and unfolded proteins accumulate within the ER lumen, leading to a cellular condition known as ER stress (Kim et al., 2018; Liu and Howell, 2010).

To restore ER homeostasis and promote cell survival, plants activate the Unfolded Protein Response (UPR), a conserved signaling network that enhances ER folding capacity, stimulates protein quality control system, and adjusts protein synthesis rates (Beaugelin et al., 2020). In *Arabidopsis thaliana*, the UPR is mediated by two major signaling branches. The first branch relies on INOSITOL-REQUIRING ENZYME 1 (IRE1) proteins and the downstream transcription factor (TF) bZIP60, whereas the second involves the ER-localized membrane TF bZIP17 and bZIP28. Activation of UPR signaling results in the transcriptional induction of genes encoding ER chaperones, folding enzymes, and components of ER-associated degradation (ERAD), thereby increasing the capacity of the ER to cope with protein folding stress (Ko et al., 2026).

IRE1 is an ER-resident transmembrane kinase/endoribonuclease that exists in three isoforms in *Arabidopsis*: IRE1A, IRE1B, and IRE1C. Among these are IRE1A and IRE1B that are the main regulators of UPR signaling in vegetative tissues. Upon ER stress perception, IRE1 proteins are supposed to homodimerize and activate their RNase domain, which mediates the unconventional splicing of *bZIP60* mRNA. This processing generates the active form of the bZIP60 TF, which subsequently regulates the expression of UPR-responsive genes (Nagashima et al., 2011). In addition to *bZIP60* splicing, IRE1 contributes to ER stress adaptation through regulated IRE1-dependent decay (RIDD), a selective mRNA degradation mechanism targeting transcripts mainly encoding secretory pathway proteins (Mishiba et al., 2013). By indirectly reducing the influx of newly synthesized proteins into the ER, RIDD contributes to restoring ER proteostasis during stress conditions (Hollien and Weissman, 2006; Kimmig et al., 2012; Mishiba et al., 2013).

The second UPR branch involves the ER membrane-bound TF bZIP17 and bZIP28. Under ER stress conditions, these proteins are transported from the ER to the Golgi apparatus, where they undergo regulated intramembrane proteolysis. The released cytosolic domains subsequently migrate to the nucleus and activate the expression of UPR-responsive genes (Liu et al., 2007).

Among the environmental challenges requiring rapid protein synthesis and extensive cellular reprogramming, pathogen attack represents a major trigger of ER stress. Plants rely on a sophisticated immune system to detect and respond to microbial threats. Recognition of pathogen- or microbe-associated molecular patterns (PAMPs/MAMPs), such as the bacterial flagellin-derived peptide flg22, activates pattern-triggered immunity (PTI), whereas recognition of pathogen-derived effector proteins initiates effector-triggered immunity (ETI) (Chisholm et al., 2006). These immune responses involve extensive transcriptional and metabolic reprogramming, including the production of defense-related proteins. Hallmarks of activated PTI include rapid production of reactive oxygen species (ROS) and deposition of callose at the cell wall. Although historically considered distinct pathways, PTI and ETI are now recognized as interconnected immune layers that share signaling components and mutually potentiate each other to establish effective plant resistance (Ngou et al., 2021).The phytohormone salicylic acid (SA) is a central regulator of plant immunity, acting as a key signaling molecule in both local defense responses (PTI and ETI) and systemic acquired resistance (Wani et al., 2017). Increasing evidence suggests that SA signaling and ER stress responses are closely interconnected (Meng et al., 2026; Poór et al., 2019). For instance, exogenous SA application induces the expression of UPR-related genes (Nagashima et al. 2014), suggesting that SA signaling can modulate ER stress responses. Conversely, ER stress conditions can stimulate SA accumulation (Wang et al., 2019), indicating reciprocal regulation between these two pathways. Nevertheless, the direct experimental evidence of signaling cross-talk between SA and acute ER stress is still lacking.

Most studies investigating ER stress responses in plants and animals rely on pharmacological induction using tunicamycin, an inhibitor of N-glycosylation, or dithiothreitol, a reducing agent that disrupts disulfide bond formation and protein folding (Chakraborty et al., 2017; Iwata and Koizumi, 2005; Wang et al., 2019, 2020). Although these approaches efficiently induce ER stress associated with impaired protein maturation, they may not fully reproduce the specific characteristics of ER stress occurring during environmental stress or pathogen infection. By assessing defense-related phenotypes and response to tunicamycin-induced ER stress in mutants impaired in SA and UPR pathways, we demonstrated the functional interplay between UPR activation and SA-mediated immune signaling, providing new insights into the coordination of ER homeostasis and plant defense responses.

## 2. Materials and methods

### Plant genotypes and growth conditions

The ecotype Columbia-0 of *Arabidopsis thaliana* was used as the wild-type (WT) genetic background. Specific UPR branches were studied using a set of double T-DNA mutants: *ire1a-2 ire1b-2* (SALK_018112, SAIL_238_F07; (Deng et al., 2013)), *bzip28-2 bzip60-1* (SALK_132285, SALK_050204; (Bao et al., 2019)), *bzip17-1 bzip60-1* (SALK_104326, SALK_050204; (Bao et al., 2019)), *bzip17-2 bzip28-2* (SALK_004048, SALK_132285; (Kim et al., 2018)) and *bzip17-2 bzip28-3* (SALK_004048, SALK_123659; (Kim et al., 2018)). The role of endogenous SA level was studied using a collection of mutants with different basal SA levels (Pluhařová et al., 2019), including the SA-deficient genotypes *pad4* (CS3806; (Zipfel et al., 2004)), *NahG* (CS1092; (Nawrath and Métraux, 1999)) and *sid2* (SALK_042603; (Wildermuth et al., 2001)), as well as the SA over-accumulators *bon1-1* (SALK_123132; (Li et al., 2007)), *snc1-1* (SALK_047058; (Zhang et al., 2003)), *bon1-1 snc1-11* (SALK_123132, SALK_047058; (Li et al., 2007)) and *pi4kβ1 pi4kβ2* (SALK_098069C, SALK_040479; (Starodubtseva et al., 2022)).

Seeds were surface sterilized with a 1.5% (w/v) solution of sodium hypochlorite (SAVO Original, Unilever, The Czech Republic) containing 0,02 % (v/v) Tween20 (Sigma Aldrich, St. Louis, Missouri, USA). Sterilized seeds were washed twice with sterile distilled water and kept in the dark at 4 °C for two days to synchronize germination. Seeds were sown in commercial substrate Jiffy-7 (Jiffy Products International AS, Norway). Plants were grown under short day conditions (14h night / 10h day with a light intensity of 100-130 μE m^−2^ s^−1^) at 22 °C, with 70 % relative humidity, in the cultivation box Snijders Labs Climate Chamber MCA1600E (Snijders Scientific, The Netherlands) and watered with fertilizer-free distilled water. Seven days after germination seedlings were transplanted into individual pots and cultivated for 4-5 weeks.

### Chemical treatments

Rosettes were sprayed with 900 µM benzothiadiazole (BTH; Bion, Syngenta, USA) or three mature fully expanded leaves (leaves 8, 9 and 10; as described by Farmer et al., 2013) were infiltrated by needleless syringe with 100 nM flg22 (EZBiolab, USA) or 0.5 µg/mL tunicamycin from *Streptomyces sp*. (T7765, Sigma-Aldrich, Germany). Mock treatments with distilled water were used as control.

### Pathogen assay

The immune response was tested with the hemibiotrophic bacterial phytopathogen *Pseudomonas syringae* pv. *tomato* (Pst) strain DC3000 (Cuppels, 1986) and its type III secretion system-deficient mutant hrcC-, which cannot inject effectors to the host plant cell (Collmer et al., 2002; Hauck et al., 2003). Bacteria was cultivated at 26°C on Pseudomonas agar medium (CN Agar, M085, HiMedia Laboratories, India) supplemented with 50µg/mL of rifampicin. Overnight bacterial culture was resuspended in a 10 mM sterile MgCl_2_ aqueous solution until OD_600_ reached 0,0005 – 0,001. This suspension was used for inoculating three mature leaves per plants by needleless syringe infiltration. Three days after inoculation, 3 leaf discs (6 mm diameter) were collected from treated leaves of each plant into 1.5 mL tubes with 1 g of ceramic beads and 1 mL of 10 mM MgCl_2_ aqueous solution. Tissues were homogenized in the FastPrep-24 instrument (MC Biomedicals, USA) two times by shaking for 40 s at a rate of 4.5m/s. Homogenized samples were serially diluted six-times with a ten-fold dilution step. Drops of diluted samples (5 μL) were deposited and left drying on square Petri dishes containing Pseudomonas agar medium containing 50 µg/mL rifampicin. Bacterial colonies (CFU, colony forming units) were counted two days later and bacterial proliferation was expressed as log_10_(CFU/mm^2^).

### Damage score

Leaves treated with tunicamycin were harvested four days after infiltration, scanned with Epson Photo V800 scanner at 600 dpi resolution and assigned a numeric value of the level of damage in accordance with the custom damage-score scale (Fig. S1).

### ROS kinetics measurement by luminol-based assay

The rapid oxidative burst caused by perception of bacterial elicitor was measured by a luminol-peroxidase assay (Trujillo, 2016; Janda et al., 2019). Leaf discs (4 mm diameter) were incubated in distilled water in the dark at room temperature for 24 h, then the water was replaced with 200 µL of reaction mixture containing 17 µg/mL luminol (Fluka, Switzerland), 10 µg/mL horseradish peroxidase (Sigma-Aldrich, Germany) and 100 nM flg22 (peptide flg22 derived from the flagellin N-terminus of *Pseudomonas* sp. (QRLSTGSRINSAKDDAAGLQIA, Tebu-Bio, France). Luminescence was recorded every 2 min with a Tecan Infinite M Plex luminometer over a 20 min kinetic interval with 3.5 s integration time per well.

### Callose deposition

To assess callose deposition leaves were syringe-infiltrated with 100 nM flg22 and, 24 h later, decolorized in ethanol/glacial acetic acid (3:1, v/v). After progressive rehydratation (2 h incubation in 60% - 40% - 20% ethanol v/v) and staining for 4 h with 0.01% aniline blue in 150 mM K_2_HPO_4_, pH 9.5 (Kalachova et al., 2019). Callose was visualized by fluorescence microscopy (Zeiss AxioImager ApoTome2, 5x objective) and quantified in Fiji/ImageJ by counting aniline blue-positive spots within regions of interest of consistent size (Schindelin et al., 2012). At least three images per sample and five non-overlapping regions of interest per image were analyzed.

### Salicylic acid quantification

SA content in tissues was measured by bioluminescence assay with *Acinetobacter sp*. ADPWH_lux biosensor (DeFraia et al., 2008). Leaves treated with tunicamycin were collected 4 days after infiltration. Plant material was weighted 80 mg minimal fresh weight per sample, 1-2 plants for one sample), and immediately flash frozen in liquid nitrogen. Tissues were homogenized with a FastPrep-24 instrument for 60 s at 4 m/s, frozen in liquid nitrogen and then mixed with 0.1 M acetate buffer, pH = 5,6 (1 weight / 2.5 volume) and homogenized again. Samples were incubated for 5 minutes on ice and centrifuged for 10 minutes at 15,000 x g at 4 °C, and the supernatant was kept for the assay. *Acinetobacter sp*. ADPWH_lux bacterial culture was incubated in liquid LB medium at 37 °C under constant shaking 120 rpm for 24 h and then passaged into fresh liquid LB medium 1:10 (v/v) and incubated for 90 min under the same conditions until OD_600_ reached 0.4. Twenty microliters of sample or standard were applied on the black 96-well plate in two technical replicates with 60 µL of liquid LB and 50 µL of bacterial culture. The plate was incubated for 1 hour at 37 °C. The luminescence was measured using Tecan Infinite M Plex (Austria). The calibration curve was prepared using a 10 ng/μL SA solution in 0.1 M acetate buffer, pH5.6 and homogenized plant extract from untreated WT plants. Final calibration points were 7.69 ng/ml, 15.38 mg/ml, 38.46 ng/ml, 76.92 ng/ml, 153.85 ng/ml, 230.77 ng/ml, 307.69 ng/ml, 384.62 ng/ml. Calibration curve was performed for each plate measurements, and each sample was assessed in 3 biological (independent plants) and 3 technical replicates (independent wells).

### Gene expression measurement by RT-qPCR

Harvested leaves were immediately frozen in liquid nitrogen and stored at -80°C. For each sample, 2-3 leaves per genotype were collected. Total RNA was isolated from leaf tissues with the Spectrum™ Plant Total RNA Kit (STRN250-1KT, Sigma-Aldrich, Merck, Germany) and treated with DNase (AM2222, DNA-free Kit, Ambion, USA), according to the manufacturers’ instructions. One microgram of RNA was reverse-transcribed with M-MLV H-Point Mutant reverse transcriptase (M3681, Promega Corporation, USA). Quantitative PCR was performed on a LightCycler 480 (Roche Diagnostics, Basel, Switzerland) with an initial denaturation at 95 °C for 5 min followed by 40 cycles of 95 °C for 30 s, 55 °C for 30 s and 72 °C for 3 min. Expression was normalized to three reference genes (*ACT2, EF1a, RHIP*) and calculated taking into account primer efficiencies (Ducloy et al., 2024). Primers are listed in Supplementary Table T1.

### Statistical analysis

Values from each experiment were processed with MS Excel and sorted data were analyzed in GraphPad Prism 8.0.1 (GraphPad Software, San Diego, CA, USA). Data were compared using one-way ANOVA and two-way ANOVA with specific post-hoc tests. Significant differences were marked with different letters or asterisks (p < 0,05). All experiments were performed in at least 3 independent biological replicates. The exact number of replicates, technical repetitions, and the statistical tests applied are stated in the figure legends.

## 3. Results

### UPR-deficient mutants show enhanced PAMP-triggered immunity in response to *Pseudomonas syringae* pv *tomato*

To test the interplay between UPR and immunity, a set of double mutants defective in either of the two UPR branches were inoculated with the strain DC3000 of *Pseudomonas syringae* pv. *tomato* (*Pst*), at low inoculum titer (OD_600_=0.001). Disease intensity was then quantified by determining the bacterial leaf content at 3 days post-inoculation (dpi, Fig. 1). Mutants *ire1a-2 ire1b-2, bzip17-1 bzip60-1* and *bzip17-2 bzip28-2* showed enhanced resistance characterized by a ten-fold decrease of the bacterial leaf concentration when compared to the WT. By contrast, *bzip28-2 bzip60-1* had a disease intensity statistically comparable to WT (Fig. 1A). The *hrcC*- strain of *Pst*, which cannot inject type III effectors, was then used to evaluate the ability of UPR-deficient mutant to mount PTI. It was inoculated at the same titer as for the previous set of experiments and the plant disease severity was also evaluated at 3 dpi. All UPR-deficient mutants, except for *bzip17-2 bzip28-*2, showed approximately a ten-fold reduced bacterial leaf content with respect to the WT genotype (Fig. 1B), indicating that the increased resistance of UPR mutants is mediated predominantly through PTI under our experimental conditions.

**Figure 1.**
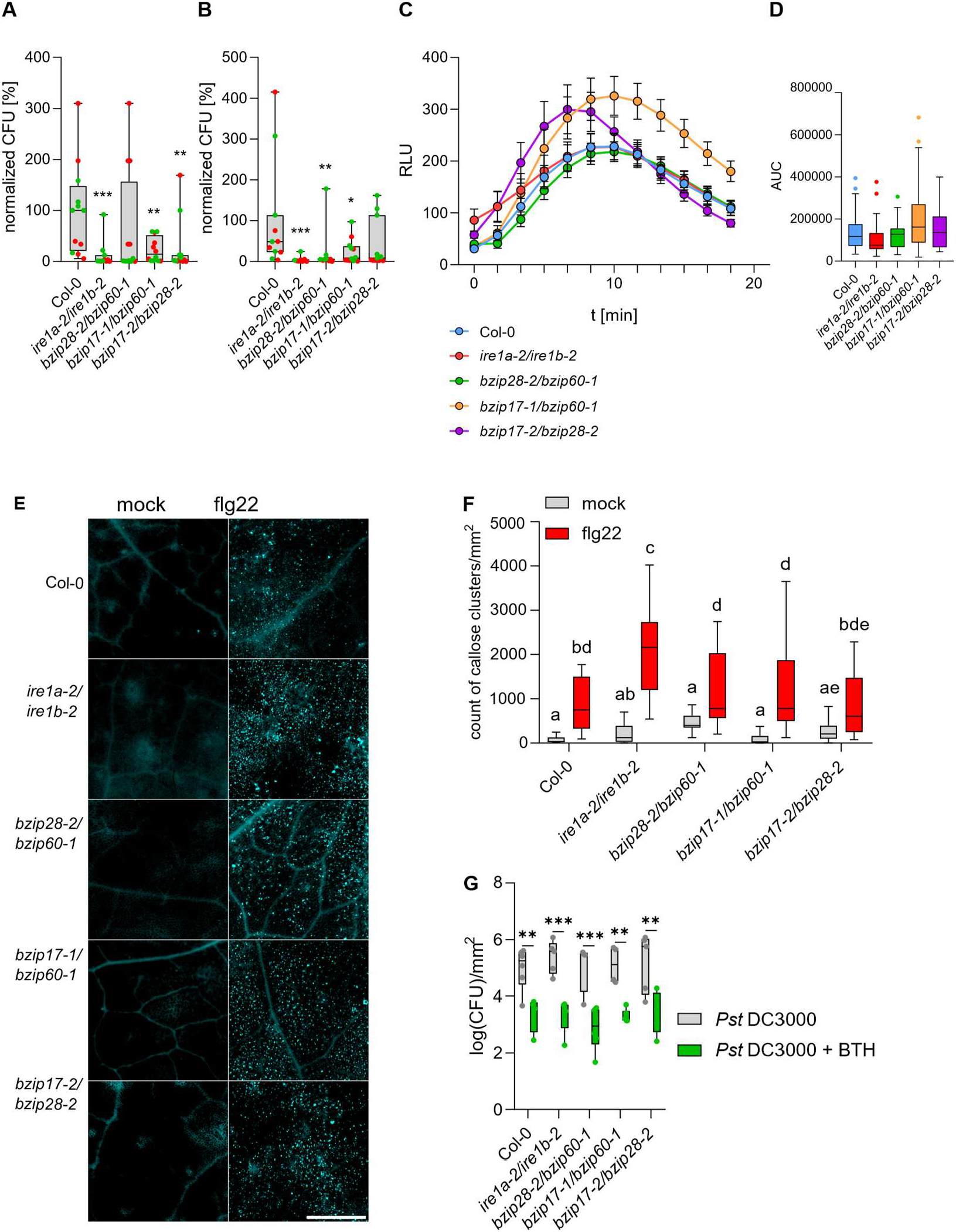
Characterization of PAMP-triggered immunity in UPR-deficient mutants. Col-0, *ire1a-2 ire1b-1, bzip28-2 bzip60-1, bzip17-1 bzip60-1* and *bzip17-2 bzip28-2*; leaves of 4-week-old plants. (A) Propagation of Pst DC3000 (OD600 = 0.001, 3 dpi); relative CFU normalized to the Col-0 mean; each genotype compared to Col-0 by Mann-Whitney test (p ≤ 0.05, n = 11-12). (B) Propagation of Pst hrcC- (OD600 = 0.001, 3 dpi); statistics as in (A) (n = 8-12). (C) Propagation of Pst DC3000 after BTH pretreatment (0.13 mg/mL spray, 1 day before inoculation); log10(CFU)/mm2; BTH compared with mock within each genotype by two-way ANOVA (n = 4-6). (D) Representative callose images (100 nM flg22, 1 dpi; aniline blue; scale = 1000 µm). (E) Callose deposit counts per mm2; two-way ANOVA, Tukey groups (n = 15-31). (F) ROS kinetics after 100 nM flg22 (luminol assay, RLU; mean ± SD). (G) ROS burst as area under the curve; Kruskal-Wallis test (n = 18-36).

### *IRE1*-deficient mutant shows enhanced callose deposition in response to flg22

To strengthen our findings, the capability of UPR-deficient mutants to engage PTI was studied using the PAMP flg22, and two PTI hallmarks were examined: oxidative burst and callose deposition. The oxidative burst was measured by luminol assay after elicitation with 100 nM flg22. All mutants reached a ROS peak approximately 10 min after treatment, except for *bzip17-2 bzip28-2*, which peaked earlier (approximately 6.5 min; Fig; 1C). The total ROS produced over 20 min was similar between genotypes (Fig. 1D), indicative of a minor role of early oxidative burst, if any, in the enhanced PTI phenotype of UPR-defective mutants. Callose deposition was measured by aniline blue staining at 24 h after 100 nM flg22 infiltration into leaves. All tested genotypes were able to accumulate callose in response to flg22, but the double mutant *ire1a-2 ire1b-2* showed a significantly greater increase in callose foci (more than doubled) than the other genotypes, consistent with a primed basal state in *IRE1*-deficient plants (Fig. 1E, F).

### UPR-deficient mutants are able to mount BTH-induced resistance

Since SA accumulation is a late signaling event known to reinforce PTI through transcriptional activation of PR genes (Tian et al., 2025), UPR-defective mutants were checked for their ability to mount a SA-dependent defense response. It was tested using BTH, a synthetic SA analog prone to activate local and systemic resistance to biotrophic pathogens (Bektas and Eulgem, 2015a). WT and mutant plants were pretreated by leaf-spraying with a 0.9 mM BTH solution at 24 h before inoculation with *Pst* DC3000 (OD_600_=0.001). As expected, BTH-pretreated WT plants were approximately 50-fold more resistant than controls sprayed with water, when bacterial leaf titer was quantified at 3 dpi. Interestingly, all UPR mutants were likewise capable of BTH-induced resistance, showing approximately 50-fold reduced bacterial propagation at 3 dpi in comparison to mock-treated plants of the same genotype (Fig. 1G). These results suggest that UPR-deficient mutants are likely still responsive to SA, allowing not to rule out the possibility that enhanced PTI observed in mutants could be due to SA downstream effectors.

### Transcriptomic evidence that IRE1A/B loss de-represses SA and defense genes under ER stress

In our previous work (Ducloy et al., 2024), a transcriptomic dataset (E-MTAB-12284) was generated to identify genes involved in chronic, proteotoxic ER stress induced by tunicamycin (100 ng/mL for 14 days) in Arabidopsis seedlings grown *in vitro*, and to distinguish which of those genes are under the control of *IRE1A* and *IRE1B* by using a double mutant (distinct from the one used in this study: SALK_002316, GABI_638B07). We reanalyzed this dataset in the light of our present findings, applying stringent thresholds (adjusted p-value < 0.05 and |log2 fold change| > 1). In WT shoots, tunicamycin treatment alone induced 716 genes and repressed 133, but among the SA-related genes only *WRKY70* and *PATHOGENESIS-RELATED PROTIEN 2* (*PR2*) passed these thresholds (Fig. 2A), indicating that chronic ER stress by itself elicits only a limited SA transcriptional response. It is markedly different for the double mutant *ire1a-3 ire1b-1* since under tunicamycin, 1553 genes were up-regulated and 690 down-regulated relative to the WT, and the up-regulated set was strongly enriched for SA and defense genes (Fig. 2B). The SA-biosynthesis genes *SALICYLIC ACID INDUCTION DEFICIENT 2* (*SID2*)/*ISOCHORISMATE SYNTHASE 1* (*ICS1*) and *AVRPPHB SUSCEPTIBLE 3* (*PBS3*), the SA markers *PR1, PR2* and *PR5*, the PTI markers *FLG22-INDUCED RECEPTOR-LIKE KINASE 1* (*FRK1*), *WRKY30* and *NDR1/HIN1-LIKE 10* (*NHL10*), and the UPR marker BINDING-IMMUNOGLOBULIN PROTEIN 3 (*BIP3*) were all significantly de-repressed (Fig. 2B, C). Gene Ontology analysis of the genes significantly up-regulated in the double mutant compared to WT under stressful conditions confirmed enrichment for response to SA, systemic acquired resistance, defense response to bacterium and fungus, and defense-related programmed cell death (Fig. 2D). Thus, although obtained with a chronic ER-stress regime applied to seedlings, this independent dataset corroborates our hypothesis that IRE1 signaling restrains SA-mediated defense and cell-death programs during ER stress.

**Figure 2.**
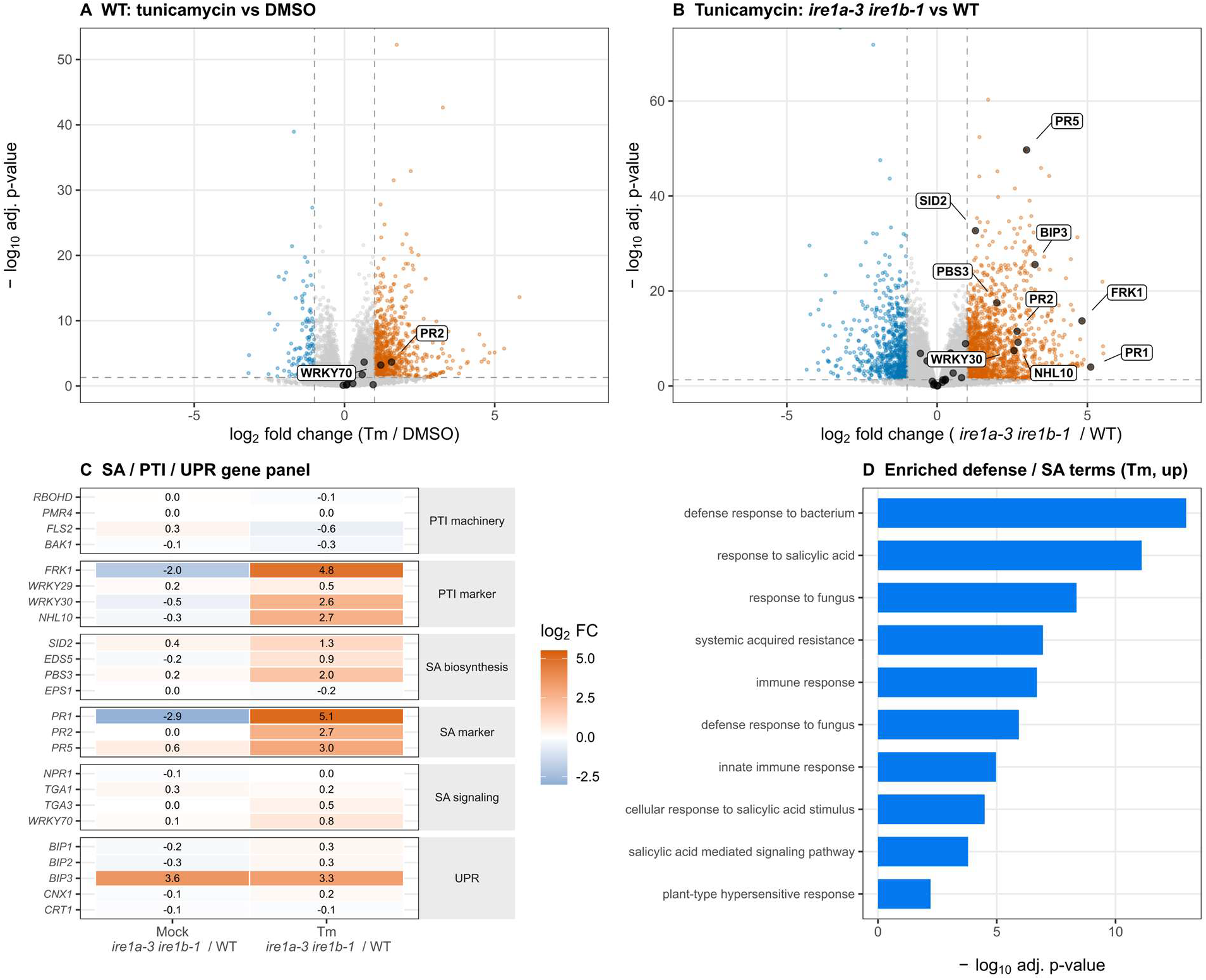
Independent transcriptomic evidence that IRE1 loss de-represses SA and defense genes under ER stress (reanalysis of Ducloy et al., 2024; E-MTAB-12284). RNA-seq of WT and *ire1a ire1b* (*ire1dm*) seedlings (shoots) under mock (DMSO) and tunicamycin (Tm), 5 replicates. (A) Volcano plot of the WT tunicamycin-vs-DMSO contrast; dashed lines mark the significance thresholds (adjusted p-value < 0.05 and |log2 fold change| > 1); major salicylic-acid-related genes (SA biosynthesis, signaling and markers) passing these thresholds are highlighted. (B) Volcano plot of the *ire1a-3 ire1b-1* / WT contrast in Tm-treated shoots; dashed lines mark the significance thresholds (adjusted p-value < 0.05 and |log2 fold change| > 1); SA/PTI/UPR panel genes passing these thresholds are highlighted (standard gene names). (C) Heatmap of log2 fold change (*ire1a-3 ire1b-1*/WT) for a curated SA-biosynthesis, SA-signaling, SA-marker, PTI-marker, PTI-machinery and UPR gene panel, under mock and Tm. (D) Gene Ontology biological-process terms enriched among genes up-regulated in *ire1a-3 ire1b-1 vs* WT under Tm.

### Acute ER stress causes SA accumulation correlated with necrotic symptoms

To keep on investigating UPR and SA interplay, the hypothesis that acute proteotoxic ER stress could promote the accumulation of SA was challenged by infiltrating tunicamycin at a high concentration of 0,5µg/mL into adult Arabidopsis leaves. As previously reported (Cai et al., 2018; Chen and Brandizzi, 2012; Munch et al., 2014; Wang et al., 2007), one such infiltration produced visible leaf damages characterized for WT plants by chlorotic necrosis, appearing within four days under our experimental design (Fig. 3A). Symptoms were scored using the damage scale described in the section Material & Methods (Fig. S1), and relative abundance of leaf SA was indirectly assessed at 4 day-post infiltration using the *Acinetobacter sp*. ADPWH_lux biosensor. Consistent with their tunicamycin hypersensitivity already reported *in vitro* (Deng et al., 2013; Kim et al., 2018; Nagashima et al., 2011), *IRE1*-deficient plants and the double mutants *bzip28-2 bzip60-1* showed the most severe symptoms induced by tunicamycin (score around 10) when compared to the WT and other mutant combinations (whose score averages 5; Fig. 3A, B). Those phenotypes were accompanied by a ten-fold and seven-fold increase in SA levels in treated *ire1a-2 ire1b-2* and in *bzip28-2 bzip60-1* mutants, respectively. No difference in basal SA levels could be recorded, irrespective of the genotypes (Fig. 3C). Taken together, these data raise the question as to whether ER-stress-induced cell death (ERSID) could be dependent on SA.

**Figure 3.**
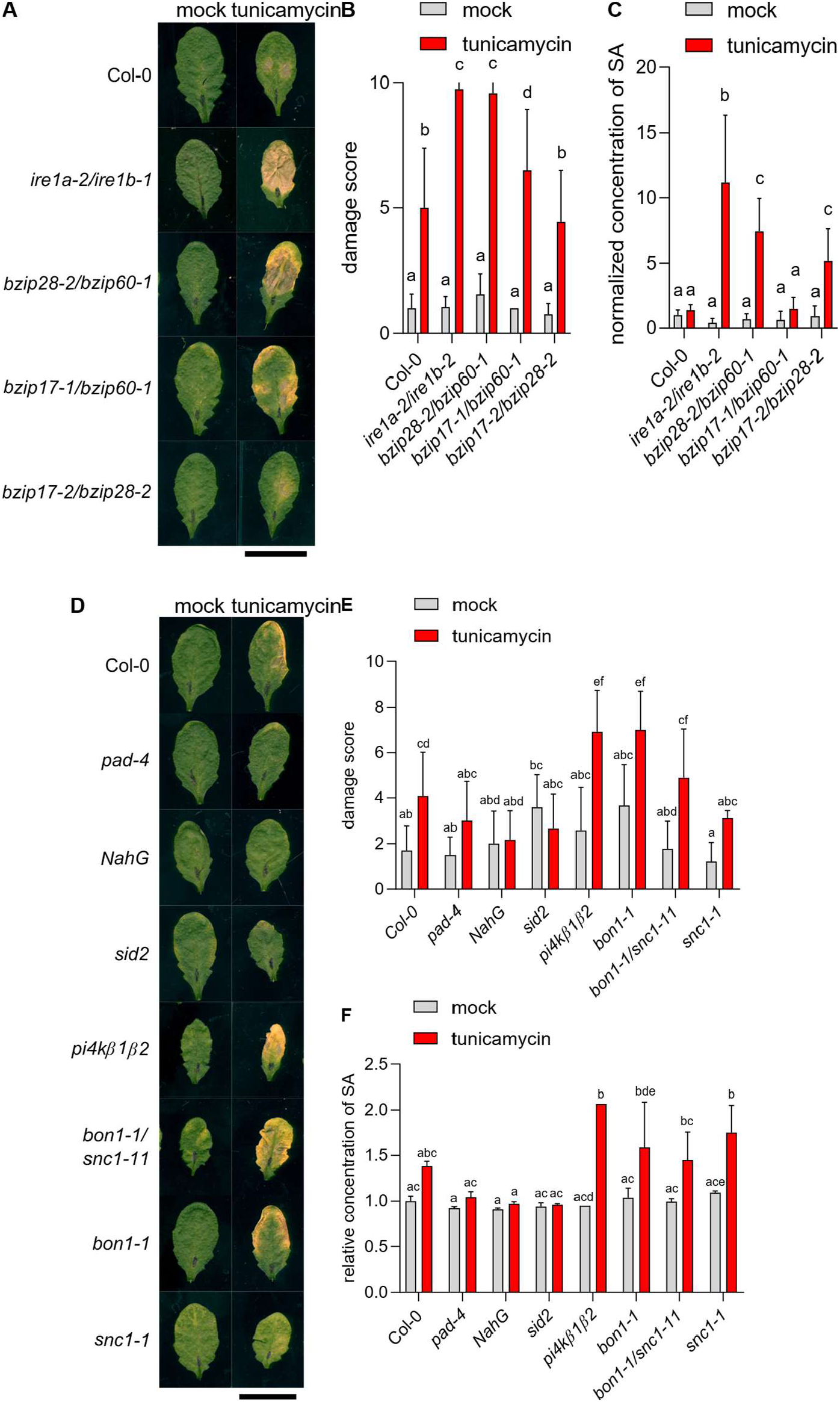
Response to acute ER stress in UPR and SA mutants. Leaves of four-week-old plants were infiltrated with 0.5 µg/mL tunicamycin or DMSO (mock), scored at 4 dpi. (A) Representative damage in UPR mutants (scale = 1 cm). (B) Damage score in UPR mutants (mean + SEM; two-way ANOVA, Tukey groups; n = 18-37). (C) Relative SA level in UPR mutants (Acinetobacter sp. ADPWH_lux biosensor; normalized to mock Col-0; two-way ANOVA; n = 6-14). (D) Representative damage in SA mutants (scale = 1 cm). (E) Damage score in SA mutants (two-way ANOVA; n = 9-12). (F) Relative SA level in SA mutants (two-way ANOVA; n = 3-4).

### ER stress-induced cell death triggered by tunicamycin is dependent on endogenous level of SA

Testing the hypothesis that endogenous SA levels could dictate cell fate under acute ER stress conditions, we have selected a set of mutants depleted in SA content and/or altered in SA signaling (*NahG* mutant, expressing salicylate hydroxylase and unable to accumulate SA (Delaney et al., 1994), *sid2-3* mutant, deficient in ISOCHORISMATE SYNTHASE 1, a key enzyme of pathogen-induced SA biosynthesis in Arabidopsis (Wildermuth et al., 2001), and *pad4* mutant, deficient in *PHENYLALANINE DEFICIENT 3* gene, an important component of SA downstream signaling and self-amplifying biosynthetic loop, Zipfel et al., 2004), together with mutants constitutively overaccumulating SA (a mutant compromised in vesicular trafficking *pi4kβ1β2* (Šašek et al., 2014), mutants with constantly de-repressed immunity *bon1-1, snc1-1* (Li et al., 2007), and the double mutant *bon1-1 snc1-11* with reverted phenotype (Pluhařová et al., 2019). Plants were leaf-challenged with high concentration of tunicamycin (0.5 mg/mL, Fig. 3D), and both the induced symptoms and relative endogenous SA content were quantified at 4 day-post infiltration (Fig. 3 E,F). The selected mutants were previously characterized for their basal SA contents under different growth conditions (Pluhařová et al., 2019), so SA content after Tm treatment was normalized to untreated controls for each genotype.

Remarkably, all three SA-deficient genotypes showed no leaf symptoms in response to tunicamycin (Fig. 3D), as their damage score (around 2) was comparable to that of untreated WT leaves (Fig. 3E). In line with our hypothesis, this was correlated with an absence of SA accumulation in the three mutant genetic backgrounds whereas tunicamycin promoted a faint, but significant increase in SA in WT plants (1.5-fold, Fig. 3F). In addition, mutants *pi4kβ1β2, snc1-1* and *bon1-1*, showing the highest monitored increased SA levels (Fig. 3F), displayed more symptoms than the tunicamycin-treated WT, with a two-point increase of their damage score with respect to WT plants (Fig. 3E). Finally, double mutant *bon1-1 snc1-11* showed symptom severity comparable to WT (Fig 3 E,F). Therefore, increased SA content seems to correlate with, and be required for, ERSID.

## 4. Discussion

The UPR is classically viewed as an adaptive mechanism that preserves ER homeostasis during proteotoxic stress, and can engage programmed cell death when ER stress cannot be resolved (Cacas, 2015; Liu and Howell, 2010; Mishiba et al., 2013) A large body of findings also accumulated that shed light on the major contribution of UPR to plant immunity through mechanisms that remain incompletely understood (Marzari et al., 2025; Poór et al., 2019). In the present study, we demonstrate that impairment of the canonical IRE1– bZIP60 branch enhances resistance to *Pst*, primarily through potentiation of PTI (Fig. 1). Our results further suggest that IRE1 signaling normally restrains excessive activation of SA-dependent defense pathways (Fig. 2) and identify SA as a major determinant of cell death during acute ER stress (Fig. 3).

To investigate the contribution of the UPR to plant immunity, we compared the growth of *Pst* DC3000 and the type III secretion-deficient mutant *Pst* hrcC^−^ in a collection of UPR-deficient mutants. Unexpectedly, mutants impaired in the IRE1–bZIP60 branch consistently exhibited enhanced resistance to both bacterial strains compared with WT plants (Fig. 1 A, B). This observation contrasts with previous reports pointing out UPR as a positive regulator of Arabidopsis immunity in response to bacterial pathogens, and, to some extent, in response to phytopathogenic fungi and oomycetes (Blanchard et al., 2025; Marzari et al., 2025; Qiang et al., 2021; Samperna et al., 2021; Xu et al., 2019). Although pathogen nature and lifestyle may explain one such discrepancy, we were concerned with the study of Chakraborty et al. (2020) showing no differences in leaf bacterial growth between WT plants and the *ire1a-2 ire1b-1* double mutant upon infection with *Pst*. A notable methodological difference between our study and that of colleagues was the bacterial inoculum used (for both DC3000 and hrcC^−^ strains). Chakraborty et al. (2020) inoculated plants with a lower bacterial concentration, which may have limited the sensitivity of the assay. We tested this hypothesis experimentally and confirmed it, as inoculation with *Pst* DC3000 at OD_600_ = 0.0005 (titer used by Chakraborty et al. 2020) failed to reveal significant differences between genotypes under our experimental conditions (Fig. S2A, B), whereas increasing the inoculum to OD_600_ = 0.001 consistently discriminated the resistance phenotypes of the UPR-deficient mutants (Fig. S2).

The enhanced resistance of UPR-deficient mutants to both *Pst* DC3000 and hrcC^−^ strains (Fig. 1A, B) indicates that loss of UPR signaling does not compromise antibacterial immunity. The reduced leaf growth of the hrcC^−^ mutant strongly argue in favor of an enhanced PTI rather than an altered recognition of bacterial effectors. This conclusion is reinforced by the increased flg22-induced callose deposition observed in the double *ire1a-2 ire1b-2* mutant (Fig. 1D, E). The early oxidative burst being largely unaffected in this mutant (Fig. 1F, G), it is possible that loss of IRE1 signaling selectively potentiates downstream PTI outputs rather than globally enhance all defense responses. This conclusion, consistent with the PTI being composed of multiple partially independent signaling modules, would need to be further investigated. This could be done by quantifying defense gene expression by qPCR and monitoring PR secretion, as already performed by others (Chakraborti et al., 2020; Moreno et al., 2012). Such approach would also be useful to deepen the involvement of bZIP28 in PTI, as the *bzip28-2-bzip60-1* mutants show callose deposition as WT plants (Fig. 1D, E).

The exact molecular basis underlying enhanced PTI in *IRE1*-deficient plants remains to be elucidated. One possibility is that constitutive ER dysfunction in these mutants establishes a primed physiological state, allowing faster or stronger activation of defense responses upon pathogen perception. Such a model is supported by our transcriptomic re-analysis (Ducloy et al. 2024), which revealed strong de-repression of genes involved in SA biosynthesis (*ICS1/SID2* and *PBS3*), SA signaling (*PR1, PR2, PR5*) and PTI (*FRK1, NHL10* and *WRKY30*) in *ire1a-3 ire1b-1* mutant seedlings subjected to ER stress (Fig. 3B, C). Together with the enrichment of Gene Ontology categories related to systemic acquired resistance, defense against bacterial and fungal pathogens, and defense-associated programmed cell death (Fig. 3D), these data suggest that IRE1 normally acts as a negative regulator preventing excessive activation of immune transcriptional programs during ER stress.

The persistence of BTH-induced resistance in all UPR-deficient mutants (Fig. 1C) further refines our interpretation of the relationship between UPR and SA signaling. BTH is a functional analogue of SA that activates defense responses downstream of SA biosynthesis through the NPR1-dependent signaling pathway (Bektas and Eulgem, 2015b). Accordingly, exogenous BTH bypasses the requirement for endogenous SA production mediated by the ICS1–EDS5–PBS3 pathway. The ability of all UPR mutants to develop BTH-induced resistance therefore indicates that downstream SA perception and signaling remain fully functional despite impaired UPR signaling. Consequently, the enhanced PTI phenotype observed in UPR-deficient plants is unlikely to result from altered SA responsiveness. Rather, our transcriptomic analyses, showing de-repression of ICS1, PBS3 and several SA-responsive genes in *ire1* double mutants under ER stress, suggest that the IRE1 branch of the UPR primarily regulates endogenous SA accumulation and/or its transcriptional activation rather than the downstream signaling machinery. This interpretation reconciles our observations with previous studies reporting altered SA responses in *ire1* mutants during ER stress (Moreno et al., 2012), as the requirement for IRE1A and IRE1B may specifically arise when endogenous SA biosynthesis is activated under conditions of ER proteotoxic stress.

Although impairment of IRE1 signaling enhanced antibacterial immunity, it markedly increased susceptibility to acute ER stress induced by tunicamycin and accelerate ERSID outcomes (Fig. 2 A, D). Both *ire1a-2 ire1b-2* and *bzip28-2 bzip60-1* plants developed severe chlorotic and necrotic foliar lesions in response to tunicamycin treatment (Fig. 2A, B), confirming previous *in vitro* studies showing that the two major UPR branches function cooperatively to preserve ER homeostasis (Deng et al., 2013; Kim et al., 2018; Mishiba et al., 2013). Notably, only mild changes non-significant in chaperone gene transcription were detected by the time of ERSID (Fig. S3). A major finding of our work is the demonstration that SA is a likely key determinant of ERSID. As reported by Wang et al. (2019), tunicamycin was found to promote SA accumulation (Fig. 2F). Remarkably, endogenous levels of SA closely correlated with symptom severity across genotypes (Fig. 2B, C, E & F). Mutants defective in SA biosynthesis or signaling (*sid2, NahG* and *pad4*) were largely protected from ERSID, whereas mutants accumulating SA exhibited enhanced tissue damage. Furthermore, SA levels near to WT in the *bon1-1 snc1-11* double mutants was associated to WT sensitivity to tunicamycin. These complementary genetic approaches provide compelling evidence that endogenous SA is not merely associated with ERSID, but is required for its full execution.

The mechanism linking ER stress to SA accumulation remains unclear. Previous studies have shown that ER stress can activate SA signaling and that SA itself can activate both the IRE1-bZIP60 and the bZIP28 arms of the UPR (Poór et al., 2019). Our transcriptomic analysis extends this concept by identifying *ICS1/SID2* and *PBS3*, two key enzymes of pathogen-induced SA biosynthesis, among the genes de-repressed in *ire1* double mutants during ER stress. These observations suggest that IRE1 signaling contributes to limiting SA production during proteotoxic stress, thereby preventing inappropriate activation of defense-associated cell death programs. Such negative regulation may represent an important mechanism allowing adaptive UPR responses to restore ER homeostasis before irreversible cell death is initiated.

Collectively, our results support a model in which IRE1 signaling fulfils two complementary functions during ER stress. Under moderate stress, IRE1 contribute to maintaining ER homeostasis while simultaneously suppressing excessive activation of SA-dependent defense pathways. Loss of IRE1 releases this repression, leading to enhanced PTI and elevated expression of defense genes, which increases resistance to bacterial pathogens. However, under severe ER stress, the inability to restore ER homeostasis results in excessive SA accumulation and activation of programmed cell death. We therefore propose that IRE1 constitutes a central molecular hub coordinating the balance between adaptation, immune activation and cell death, thereby optimizing plant fitness under fluctuating environmental conditions (Fig. 4).

**Figure 4.**
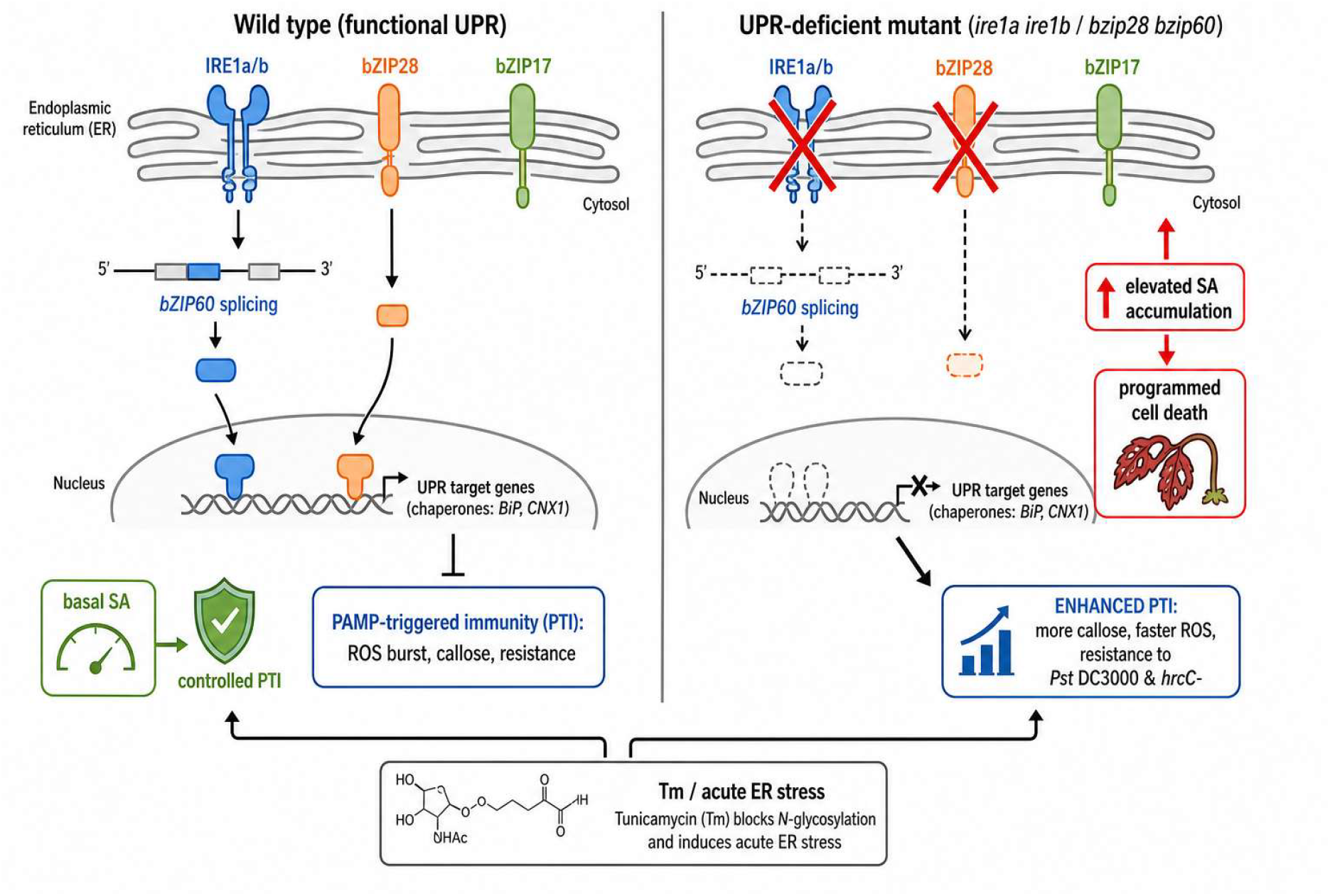
Working model: the UPR restrains PTI and buffers salicylic-acid-dependent cell death during ER stress. In wild type (left), a functional UPR (IRE1-bZIP60 and bZIP28 branches) maintains ER homeostasis and restrains PTI, keeping SA at a basal level. In UPR-deficient mutants (right), loss of IRE1/bZIP28 signaling releases the brake on PTI (enhanced callose, altered ROS burst, resistance to Pst DC3000 and hrcC-) and, under acute ER stress (tunicamycin), leads to elevated SA accumulation and SA-dependent programmed cell death.

## 5. Conclusion

We have linked UPR signaling and SA-mediated cell death, providing a new perspective on plant physiology under stress. Our experiments demonstrate that the UPR restrains PTI under non-stress conditions — so that UPR-deficient mutants are primed and more resistant to bacterial pathogens — and that acute ER stress drives salicylic acid accumulation, which in turn executes programmed cell death. By separating these two directional interactions, our work reconciles conflicting reports and identifies the IRE1–bZIP60 axis and SA as the overlapping nodes between defense and ER-stress signaling, with potential applications in resistance-inducer design and crop stress resilience.

## Author contribution

VC, TK - conceptualization; VC, AA, HL - data curation; VC, TK - formal analysis; TK, HL, JLC - methodology; TK, LB - funding acquisition, project administration; VC, TK, JLC - writing, original draft; VC, JLC, MA, AD, AA, LB, HL - review & editing. All authors commented and approved the manuscript. Authors declare no conflict of interests.

## Ethical approval statement

Authors confirm that all the research meets the ethical guidelines, including adherence to the legal requirements of the study country.

## Data availability statement

The data underlying this article will be shared on reasonable request to the corresponding author.

## Supplementary Data

**Figure S1. Tunicamycin damage scoring scale**. Representative leaves illustrating the 0-10 damage score used to quantify tunicamycin-induced leaf damage at 4 dpi.

**Figure S2. Inoculum density determines the sensitivity of the resistance assay**. Propagation of Pst DC3000 in Col-0 and UPR-deficient mutants after inoculation at OD600 = 0.0005 or OD600 = 0.001 (3 dpi). The lower inoculum did not reveal significant differences between genotypes, whereas the higher inoculum resolved the increased resistance of UPR mutants.

**Figure S3. Chaperone gene expression after tunicamycin treatment**. Relative expression of BiP1/2 (A), BiP3 (B) and CNX1 (C) in leaves of 4-week-old Col-0, ire1a-2/ire1b-1, bzip28-2/bzip60-1, bzip17-1/bzip60-1 and bzip17-2/bzip28-2 infiltrated with 0.5 µg/mL tunicamycin and harvested at 4 dpi. Expression normalized to ACT2, EF1a and RHIP and primer efficiencies; mean log2(fold change) ± SEM relative to WT control; two-way ANOVA (n = 2-3).

## References

Bao, Y., Bassham, D.C., Howell, S.H., 2019. A Functional Unfolded Protein Response Is Required for Normal Vegetative Development. Plant Physiol 179, 1834–1843. 10.1104/pp.18.01261

Beaugelin, I., Chevalier, A., D’Alessandro, S., Ksas, B., Havaux, M., 2020. Endoplasmic reticulum-mediated unfolded protein response is an integral part of singlet oxygen signalling in plants. The Plant Journal n/a. 10.1111/tpj.14700

Bektas, Y., Eulgem, T., 2015a. Synthetic plant defense elicitors. Front. Plant Sci. 5. 10.3389/fpls.2014.00804

Bektas, Y., Eulgem, T., 2015b. Synthetic plant defense elicitors. Front. Plant Sci. 5. 10.3389/fpls.2014.00804

Blanchard, C., Aimé, S., Ducloy, A., Hichami, S., Azzopardi, M., Cacas, J.-L., Lamotte, O., 2025. The IRE1-bZIP60 branch of the unfolded protein response is required for the Arabidopsis immune response to Botrytis cinerea. J Exp Bot 76, 5542–5558. 10.1093/jxb/eraf244

Cacas, J.-L., 2015. Out for a Walk Along the Secretory Pathway During Programmed Cell Death, in: Gunawardena, A.N., McCabe, P.F. (Eds.), Plant Programmed Cell Death. Springer International Publishing, Cham, pp. 123–161. 10.1007/978-3-319-21033-9_6

Cai, Y.-M., Yu, J., Ge, Y., Mironov, A., Gallois, P., 2018. Two proteases with caspase-3-like activity, cathepsin B and proteasome, antagonistically control ER-stress-induced programmed cell death in Arabidopsis. New Phytologist 218, 1143–1155. 10.1111/nph.14676

Chakraborty, R., Macoy, D.M., Lee, S.Y., Kim, W.-Y., Kim, M.G., 2017. Tunicamycin-induced endoplasmic reticulum stress suppresses plant immunity. Appl Biol Chem 60, 623–630. 10.1007/s13765-017-0319-3

Chakraborty, R., Uddin, S., Macoy, D.M., Park, S.O., Van Anh, D.T., Ryu, G.R., Kim, Y.H., Lee, J.-Y., Cha, J.-Y., Kim, W.-Y., Lee, S.Y., Kim, M.G., 2020. Inositol-requiring enzyme 1 (IRE1) plays for AvrRpt2-triggered immunity and RIN4 cleavage in Arabidopsis under endoplasmic reticulum (ER) stress. Plant Physiol Biochem 156, 105–114. 10.1016/j.plaphy.2020.09.002

Chen, Y., Brandizzi, F., 2012. AtIRE1A/AtIRE1B and AGB1 independently control two essential unfolded protein response pathways in Arabidopsis. Plant J 69, 266–277. 10.1111/j.1365-313X.2011.04788.x

Chisholm, S.T., Coaker, G., Day, B., Staskawicz, B.J., 2006. Host-Microbe Interactions: Shaping the Evolution of the Plant Immune Response. Cell 124, 803–814. 10.1016/j.cell.2006.02.008

DeFraia, C.T., Schmelz, E.A., Mou, Z., 2008. A rapid biosensor-based method for quantification of free and glucose-conjugated salicylic acid. Plant Methods 4, 28. 10.1186/1746-4811-4-28

Delaney, T.P., Uknes, S., Vernooij, B., Friedrich, L.B., Weymann, K., Negrotto, D.V., Gaffney, T., Gut-Rella, M., Kessmann, H., Ward, E., Ryals, J.A., 1994. A central role of salicylic Acid in plant disease resistance. Science 266, 1247–1250. 10.1126/science.266.5188.1247

Deng, Y., Srivastava, R., Howell, S.H., 2013. Protein kinase and ribonuclease domains of IRE1 confer stress tolerance, vegetative growth, and reproductive development in Arabidopsis. Proc Natl Acad Sci U S A 110, 19633–19638. 10.1073/pnas.1314749110

Ducloy, A., Azzopardi, M., Ivsic, C., Cueff, G., Sourdeval, D., Charif, D., Cacas, J.-L., 2024. A transcriptomic dataset for investigating the Arabidopsis Unfolded Protein Response under chronic, proteotoxic endoplasmic reticulum stress. Data Brief 53, 110243. 10.1016/j.dib.2024.110243

Hollien, J., Weissman, J.S., 2006. Decay of endoplasmic reticulum-localized mRNAs during the unfolded protein response. Science 313, 104–107. 10.1126/science.1129631

Iwata, Y., Koizumi, N., 2005. An Arabidopsis transcription factor, AtbZIP60, regulates the endoplasmic reticulum stress response in a manner unique to plants. Proceedings of the National Academy of Sciences 102, 5280–5285. 10.1073/pnas.0408941102

Kalachova, T., Leontovyčová, H., Iakovenko, O., Pospíchalová, R., Maršík, P., Klouček, P., Janda, M., Valentová, O., Kocourková, D., Martinec, J., Burketová, L., Ruelland, E., 2019. Interplay between phosphoinositides and actin cytoskeleton in the regulation of immunity related responses in Arabidopsis thaliana seedlings. Environmental and Experimental Botany 103867. 10.1016/j.envexpbot.2019.103867

Kim, J.-S., Yamaguchi-Shinozaki, K., Shinozaki, K., 2018. ER-Anchored Transcription Factors bZIP17 and bZIP28 Regulate Root Elongation. Plant Physiol 176, 2221–2230. 10.1104/pp.17.01414

Kimmig, P., Diaz, M., Zheng, J., Williams, C.C., Lang, A., Aragón, T., Li, H., Walter, P., 2012. The unfolded protein response in fission yeast modulates stability of select mRNAs to maintain protein homeostasis. Elife 1, e00048. 10.7554/eLife.00048

Ko, D.K., Danieli, R., Brandizzi, F., 2026. Rewiring the unfolded protein response for plant growth recovery after stress. Plant Communications 7, 101924. 10.1016/j.xplc.2026.101924

Li, Y., Yang, S., Yang, H., Hua, J., 2007. The TIR-NB-LRR Gene SNC1 Is Regulated at the Transcript Level by Multiple Factors. MPMI 20, 1449–1456. 10.1094/MPMI-20-11-1449

Liu, J.-X., Howell, S.H., 2010. Endoplasmic Reticulum Protein Quality Control and Its Relationship to Environmental Stress Responses in Plants. Plant Cell 22, 2930–2942. 10.1105/tpc.110.078154

Liu, J.-X., Srivastava, R., Che, P., Howell, S.H., 2007. An endoplasmic reticulum stress response in Arabidopsis is mediated by proteolytic processing and nuclear relocation of a membrane-associated transcription factor, bZIP28. Plant Cell 19, 4111–4119. 10.1105/tpc.106.050021

Marzari, T., Poilvert, E., Blanchard, C., Leborgne-Castel, N., Poinssot, B., Cacas, J.-L., Lamotte, O., Gayral, M., 2025. Endoplasmic reticulum proteostasis contributes to shape plant immunity: Perspectives from plant and pathogen sides. Current Plant Biology 44, 100552. 10.1016/j.cpb.2025.100552

Meng, Z., Zheng, S., Brandizzi, F., Liu, Y., Li, C., Ai, C., Wang, Y., Qi, J., Zhang, X., 2026. Unfolding Plant Defence: Endoplasmic Reticulum Stress Signalling at the Plant-Pathogen Interface. Plant Biotechnology Journal 24, 4799–4822. 10.1111/pbi.70662

Mishiba, K., Nagashima, Y., Suzuki, E., Hayashi, N., Ogata, Y., Shimada, Y., Koizumi, N., 2013. Defects in IRE1 enhance cell death and fail to degrade mRNAs encoding secretory pathway proteins in the Arabidopsis unfolded protein response. Proceedings of the National Academy of Sciences 110, 5713–5718. 10.1073/pnas.1219047110

Munch, D., Rodriguez, E., Bressendorff, S., Park, O.K., Hofius, D., Petersen, M., 2014. Autophagy deficiency leads to accumulation of ubiquitinated proteins, ER stress, and cell death in Arabidopsis. Autophagy 10, 1579–1587. 10.4161/auto.29406

Nagashima, Y., Mishiba, K.-I., Suzuki, E., Shimada, Y., Iwata, Y., Koizumi, N., 2011. Arabidopsis IRE1 catalyses unconventional splicing of bZIP60 mRNA to produce the active transcription factor. Sci Rep 1, 29. 10.1038/srep00029

Nawrath, C., Métraux, J.-P., 1999. Salicylic Acid Induction–Deficient Mutants of Arabidopsis Express PR-2 and PR-5 and Accumulate High Levels of Camalexin after Pathogen Inoculation. The Plant Cell 11, 1393–1404. 10.1105/tpc.11.8.1393

Ngou, B.P.M., Ahn, H.-K., Ding, P., Jones, J.D.G., 2021. Mutual potentiation of plant immunity by cell-surface and intracellular receptors. Nature 592, 110–115. 10.1038/s41586-021-03315-7

Pluhařová, K., Leontovyčová, H., Stoudková, V., Pospíchalová, R., Maršík, P., Klouček, P., Starodubtseva, A., Iakovenko, O., Krčková, Z., Valentová, O., Burketová, L., Janda, M., Kalachova, T., 2019. “Salicylic Acid Mutant Collection” as a Tool to Explore the Role of Salicylic Acid in Regulation of Plant Growth under a Changing Environment. International Journal of Molecular Sciences 20, 6365. 10.3390/ijms20246365

Poór, P., Czékus, Z., Tari, I., Ördög, A., 2019. The Multifaceted Roles of Plant Hormone Salicylic Acid in Endoplasmic Reticulum Stress and Unfolded Protein Response. Int J Mol Sci 20, 5842. 10.3390/ijms20235842

Qiang, X., Liu, X., Wang, X., Zheng, Q., Kang, L., Gao, X., Wei, Y., Wu, W., Zhao, H., Shan, W., 2021. Susceptibility factor RTP1 negatively regulates Phytophthora parasitica resistance via modulating UPR regulators bZIP60 and bZIP28. Plant Physiol 186, 1269–1287. 10.1093/plphys/kiab126

Samperna, S., Boari, A., Vurro, M., Salzano, A.M., Reveglia, P., Evidente, A., Gismondi, A., Canini, A., Scaloni, A., Marra, M., 2021. Arabidopsis Defense against the Pathogenic Fungus Drechslera gigantea Is Dependent on the Integrity of the Unfolded Protein Response. Biomolecules 11, 240. 10.3390/biom11020240

Šašek, V., Janda, M., Delage, E., Puyaubert, J., Guivarc’h, A., Maseda, E.L., Dobrev, P.I., Caius, J., Bóka, K., Valentová, O., Burketová, L., Zachowski, A., Ruelland, E., 2014. Constitutive salicylic acid accumulation in pi4kIIIβ1β2 Arabidopsis plants stunts rosette but not root growth. New Phytologist 203, 805–816. 10.1111/nph.12822

Schindelin, J., Arganda-Carreras, I., Frise, E., Kaynig, V., Longair, M., Pietzsch, T., Preibisch, S., Rueden, C., Saalfeld, S., Schmid, B., Tinevez, J.-Y., White, D.J., Hartenstein, V., Eliceiri, K., Tomancak, P., Cardona, A., 2012. Fiji: an open-source platform for biological-image analysis. Nat Methods 9, 676–682. 10.1038/nmeth.2019

Starodubtseva, A., Kalachova, T., Retzer, K., Jelínková, A., Dobrev, P., Lacek, J., Pospíchalová, R., Angelini, J., Guivarc’h, A., Pateyron, S., Soubigou-Taconnat, L., Burketová, L., Ruelland, E., 2022. An Arabidopsis mutant deficient in phosphatidylinositol-4-phosphate kinases ß1 and ß2 displays altered auxin-related responses in roots. Sci Rep 12, 1–16. 10.1038/s41598-022-10458-8

Tian, H., Xu, L., Li, X., Zhang, Y., 2025. Salicylic acid: The roles in plant immunity and crosstalk with other hormones. Journal of Integrative Plant Biology 67, 773–785. 10.1111/jipb.13820

Wang, S., Narendra, S., Fedoroff, N., 2007. Heterotrimeric G protein signaling in the Arabidopsis unfolded protein response. Proceedings of the National Academy of Sciences 104, 3817–3822. 10.1073/pnas.0611735104

Wang, W., Li, Xin, Zhu, M., Tang, X., Wang, Z., Guo, K., Zhou, Y., Sun, Y., Zhang, W., Li, Xiaofang, 2019. Arabidopsis GAAP1 to GAAP3 Play Redundant Role in Cell Death Inhibition by Suppressing the Upregulation of Salicylic Acid Pathway Under Endoplasmic Reticulum Stress. Front. Plant Sci. 10. 10.3389/fpls.2019.01032

Wang, Y., Zhang, L., He, Z., Deng, J., Zhang, Z., Liu, L., Ye, W., Liu, S., 2020. Tunicamycin induces ER stress and inhibits tumorigenesis of head and neck cancer cells by inhibiting N-glycosylation. Am J Transl Res 12, 541–550.

Wani, A.B., Chadar, H., Wani, A.H., Singh, S., Upadhyay, N., 2017. Salicylic acid to decrease plant stress. Environ Chem Lett 15, 101–123. 10.1007/s10311-016-0584-0

Wildermuth, M.C., Dewdney, J., Wu, G., Ausubel, F.M., 2001. Isochorismate synthase is required to synthesize salicylic acid for plant defence. Nature 414, 562–565. 10.1038/35107108

Xu, Z., Song, N., Ma, L., Wu, J., 2019. IRE1-bZIP60 Pathway Is Required for Nicotiana attenuata Resistance to Fungal Pathogen Alternaria alternata. Front. Plant Sci. 10. 10.3389/fpls.2019.00263

Zhang, Y., Goritschnig, S., Dong, X., Li, X., 2003. A Gain-of-Function Mutation in a Plant Disease Resistance Gene Leads to Constitutive Activation of Downstream Signal Transduction Pathways in suppressor of npr1-1, constitutive 1. The Plant Cell 15, 2636–2646. 10.1105/tpc.015842

Zipfel, C., Robatzek, S., Navarro, L., Oakeley, E.J., Jones, J.D.G., Felix, G., Boller, T., 2004. Bacterial disease resistance in Arabidopsis through flagellin perception. Nature 428, 764–767. 10.1038/nature02485

