## Supplementary figures and images for "Unfolded Protein Response signaling restricts PTI and limits salicylic acid-dependent cell death under ER stress"

### Supplementary fig S1

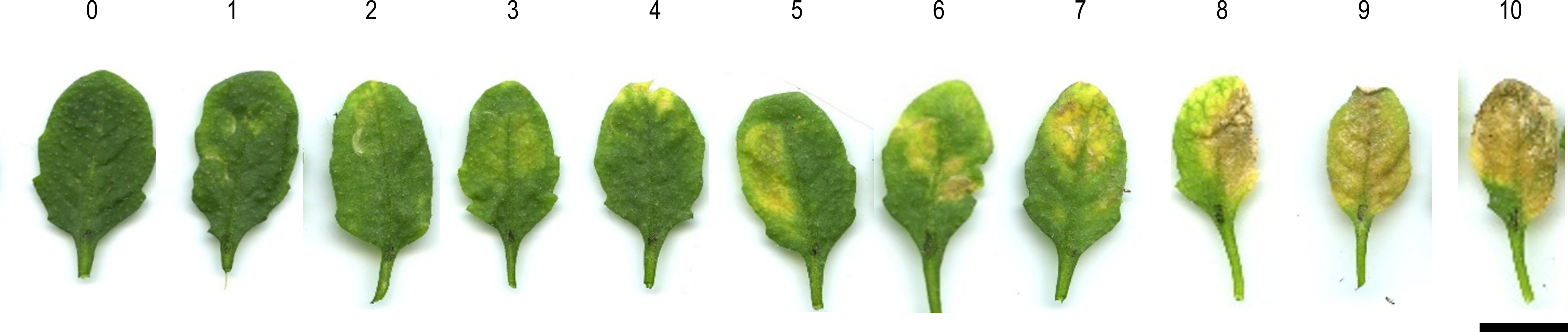

### Supplementary fig S2

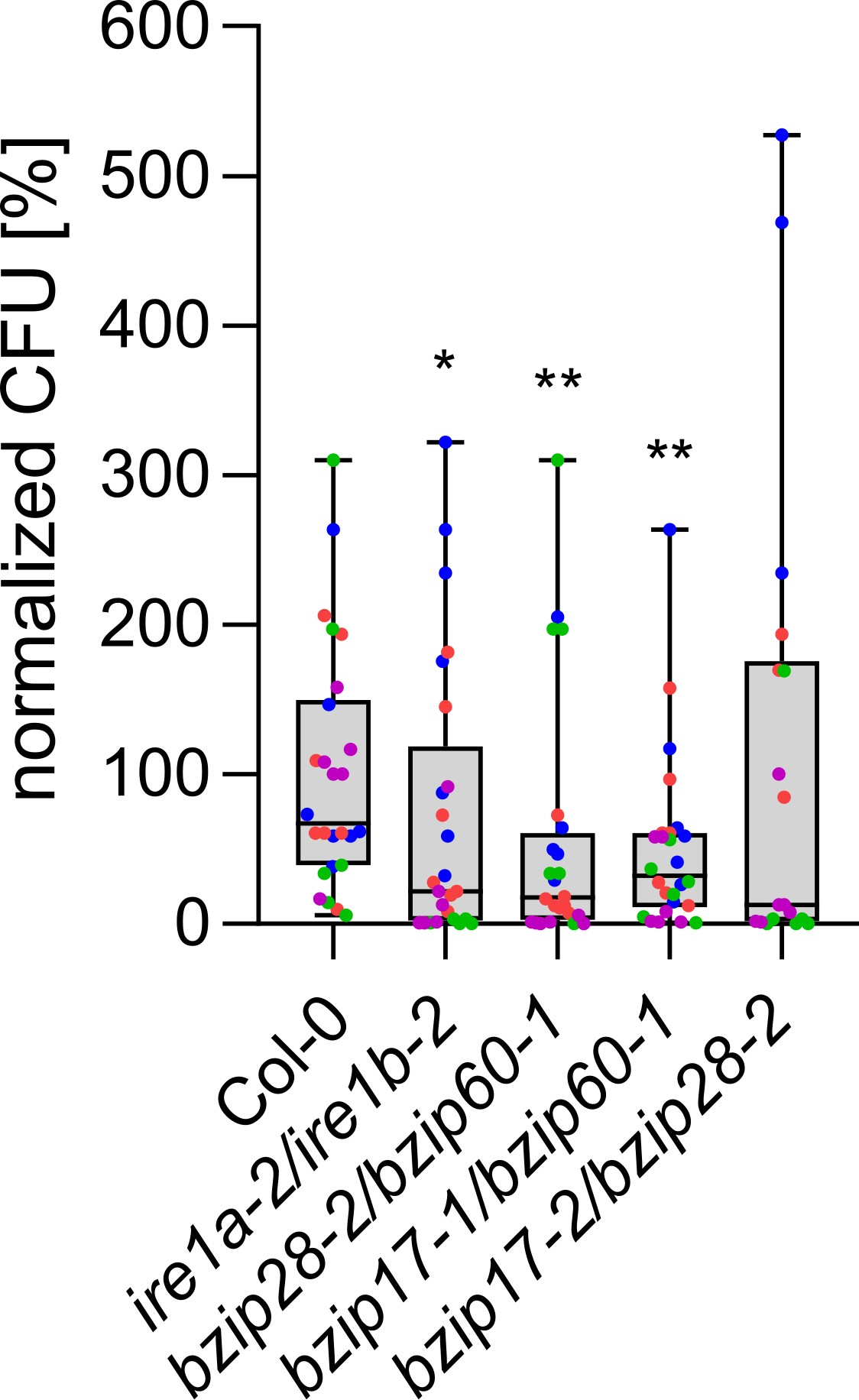

### Supplementary fig S3

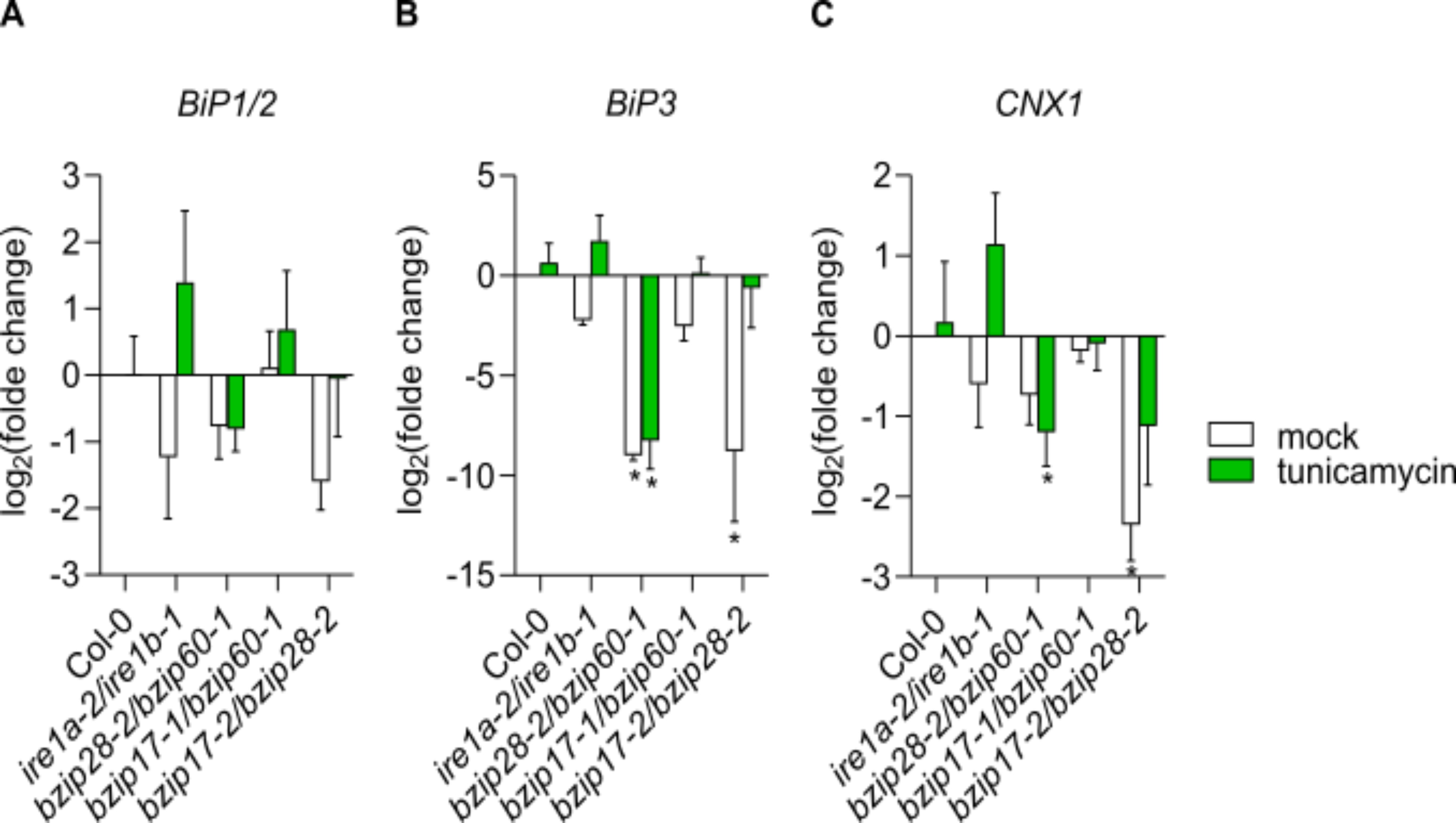
